# A Multi-lensed Comparative Analysis of Select Secondary Metabolites Produced by Kale, *Brassica oleracea*, in Simulated Microgravity Versus Gravity Conditions

**DOI:** 10.1101/2025.09.29.679299

**Authors:** Rita Dill, Yanru Li, Abdullahi Iro, George N. Ude, Supriyo Ray, Jie Yan, Anne A. Osano

## Abstract

Extended journeys through space are a goal of NASA. Yet, astronauts will face elevated health risks from microgravity and radiation as journeys continue for longer time periods. Approaches to combatting these health risks consist of growing fresh super foods in space for astronaut consumption while in flight. However, while a great deal is known about the effects of microgravity of humans, little is known about its effects on the nutrient profiles of plants. Endeavors towards understanding more about these effects are currently funded by NASA grants. Kale, a metabolite and specifically a flavonoid-rich crop, stands as a promising candidate for growth on space flights. We observed the effects of simulated microgravity broadly on the F1 cultivar, Starbor Kale metabolomics, and further focused on flavonoid content, using a 2-D clinostat. Extracts of kale were analyzed by proton nuclear magnetic resonance (^1^H NMR), and high-performance thin layer chromatography (HPTLC). ^1^H NMR spectra of clinostat-grown kale showed that samples from simulated microgravity conditions had an increased number of peaks in the aromatic region (6.5 to 8.5 ppm) when compared with gravity grown kale. HPTLC confirmed greater banding in medium- and high-polarity solvent systems, while low-polarity extracts showed no differences. Overall, we noted that the microgravity grown kale had greater amounts of bands present. These results signal that microgravity stressors may be connected to the increased secondary metabolite production in kale. Our findings underscore kale to be a prospective crop to be grown in space flight to combat effects of microgravity.

## 1. Introduction

With the goals of space flights broadening to longer durations, the monitoring of nutrition and health for astronauts becomes a heightened concern. Extensive research has shown that astronauts are at risk for a myriad of health challenges which are expected to worsen as time in space increases ^39-44^. The majority of foods consumed by astronauts are prepackaged, freeze-dried foods. Unfortunately, many of the nutrients in prepacked foods are reported to degrade over time ^45^. This proves a logistical challenge as there is a desire to increase time in spaceflight. While the current food made available to astronauts in spaceflight is invaluable, NASA has put forth efforts to grow nutritious, easily absorbed fresh foods in space with hopes that nutrients from foods may combat the health challenges in real-time while in space flight ^46-51^. Notably, these efforts include programs such as, Vegetable Production System, Passive Orbital Nutrient Delivery System, The Advanced Plant Habitat, and the XROOTS study.

Currently, there are many health challenges for astronauts when in space; it is anticipated that these challenges will be amplified when flights become longer. These challenges are connected to microgravity conditions and cosmic radiation in space. Astronauts are known to experience muscular atrophy, loss in bone density, cancer and variations in body fluid dispersion ^52-54^. These physical changes then translate to body weakness, swelling, changes in the cardiovascular system, and an increased risk of bone fractures when astronauts return to gravity conditions ^23^. While the current food sources used by astronauts are well-utilized, NASA has facilitated research grants to determine if leafy-green vegetables may be grown on spacecrafts and consumed by those in space flight. This may offer astronauts access to vital, small compounds such as flavonoids. The goal of growing, harvesting and consuming fresh food in space may lead to the achievement of longer space flights.

The ramifications of microgravity are well-investigated within human subjects ^55-57^. Yet, the effects of microgravity on the quality of food grown, and specifically the effects of microgravity general metabolomics and specifically, flavonoids requires further research as this is currently a developing field. However, some prior NASA-funded research has shown that microgravity may cause wheat plants to grow to greater heights, and that general nutrient content remains safe for human consumption ^**24**^. As there is minimal space aboard the international space station (ISS), many researchers have successfully used devices such as 2 and 3-D clinostats to simulate the effects of microgravity ^**25-26**^.

A plant that is filled with a vast array of metabolites and flavonoids is the superfood plant, kale. Kale possesses several metabolites that are useful in fighting against cancer, bone density loss and the effects of cosmic radiation ^60-62^. Kale is known to be rich in antioxidants, which, after further metabolism, can enhance levels of enzymes that perform detoxing in the body ^63-65^. These antioxidants may also assist in the prevention of DNA damage from the cosmic radiation. Kale is also a potent source of vitamin K and calcium ^59^. These two are paramount for general bone health and minimizing bone density loss ^60-62^.

While kale has many metabolites within its metabolomic profile, kale is renowned for its flavonoid content (Table 1.1) ^3^. Flavonoids constitute a large class of small compounds found in plants ^1-2^. Flavonoids are documented to have antioxidant and anti-inflammatory properties ^4-11^. A few of the modes by which flavonoids function are by reducing cellular injury and elevating natural repair systems within the body ^12-20^

**Table 1.1.**
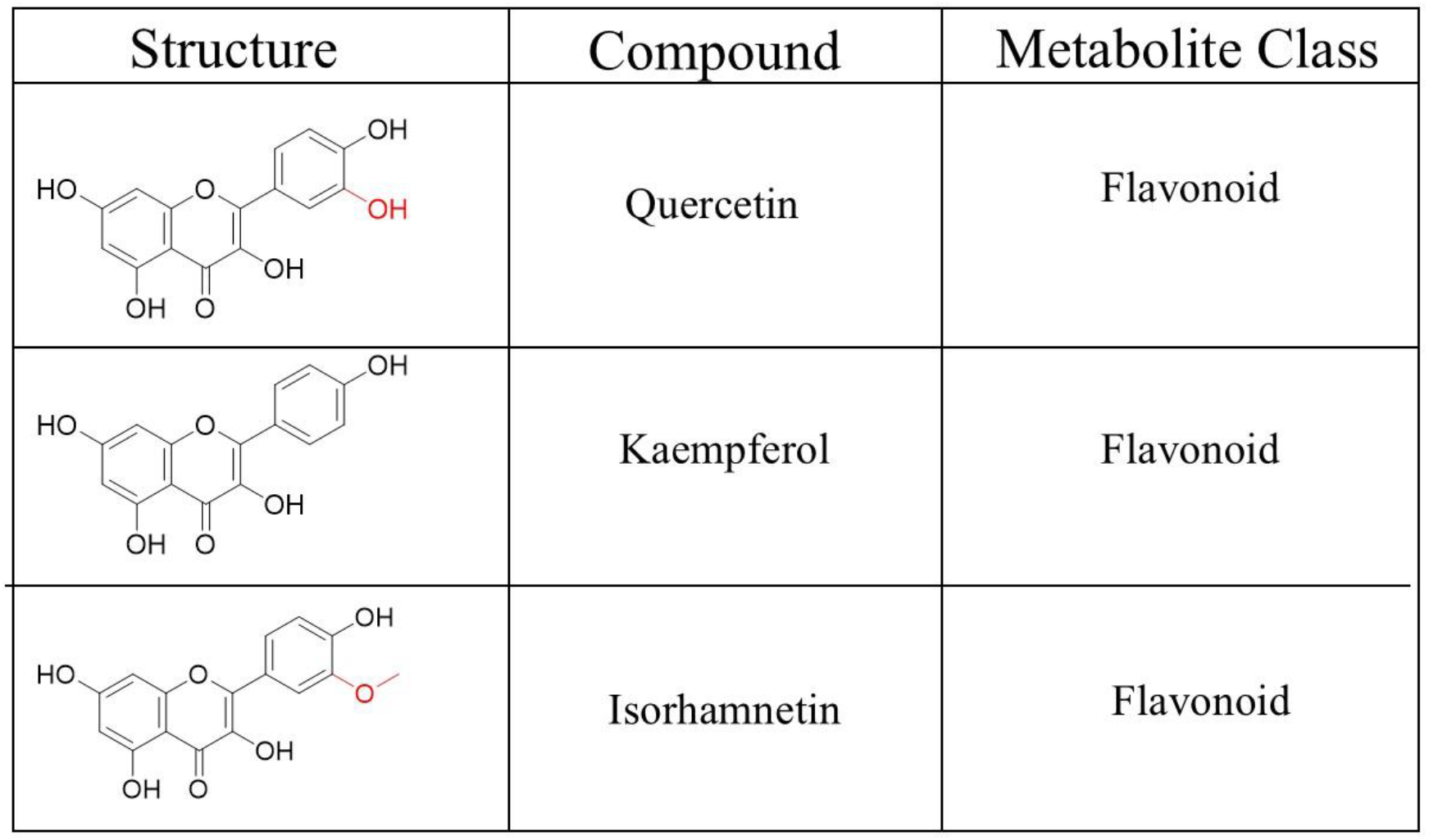
Examples of common flavonoids reported in kale ^3^ and key functional group differences are noted in red.

| Structure | Compound | Metabolite Class |
| --- | --- | --- |
|  | Quercetin | Flavonoid |
|  | Kaempferol | Flavonoid |
|  | Isorhamnetin | Flavonoid |

It is evident that kale has many metabolomic properties that nominate it as a strong candidate for astronaut food consumption research, as it may minimize the effects of microgravity and cosmic radiation. However, to perform any research studies using kale it is recommended to use an F1 cultivar crop ^58^. F1 crops tend to grow in uniform fashion, and this is ideal for the implementation of biological replicates in research. An example of an F1 kale plant cultivar is the Starbor kale.

Nuclear Magnetic Resonance (NMR) provides users with a specific understanding of the nuclei pertaining to the metabolites, and their various electronic environments ^30-32^. A rapid and simple 1-D proton NMR experiment can reveal to users the many different protons pertaining to specific functional groups or molecules, such as flavonoids. The presence of different functional groups is denoted by their different peaks; different peaks in different chemical shifts imply variations in the functional groups which constitute the metabolites present. Flavonoids are known to appear within the aromatic chemical shift range and thus, NMR is well-suited for efficient comparison of the proton NMR peaks from kale grown in different conditions. An additional planar chromatography technique, high-performance thin layer chromatography (HPTLC) has also been used to study flavonoids ^33-35^. HPTLC serves as a high throughput approach for comparing their flavonoid profiles from vegetables grown in different conditions. Combined, NMR and HPTLC provide researchers with dual methods to compare the flavonoid content found in vegetables.

Herein we report on NASA-funded research, by which we grew a leafy-green vegetable, Kale. This kale was grown in simulated microgravity environments by means of a 2-D clinostat. A control kale was grown in gravity conditions as well. The experimental and control kale were grown in soil conditions. We implemented previously published extraction protocols to target the extraction of flavonoids from the kale leaves. The extracts were subsequently studied by NMR and HPTLC analysis. The goal of our research was to compare the two conditions of gravity and simulated microgravity to test if there were notable metabolite, and more concisely flavonoid, differences between the two. We hypothesized that the experimental kale would possess different enhanced metabolite and flavonoid content. This logic was rooted in the logic that plants which undergo the stressor effects of microgravity may generate a greater number of secondary metabolites in response to such stressors. While a more extensive study on kale monitored over the course of an entire growing season is needed for future work, we believe this research will be significant in furthering the current depths of knowledge regarding the effects of simulated microgravity on flavonoid content in kale.

## 2. Materials and Methods

### 2.1. Sample preparation

Seedlings (Variety: Starbor kale) were purchased from West Coast Seeds Inc. All reagents and solvents were purchased from commercial suppliers and used without further purification. The kale seedlings were germinated using rockwool support and water for 3 weeks’ time. 1-D Proton NMR data was executed on 400 MHz Avance II spectrometer. NMR Bar chart was made using python in-house codes and all scripts are available upon request by Anne Osano. HPTLC was performed using modified previously published work ^36^. All NMR, and HPTLC data was collected at 6 weeks post planting seedlings.

### 2.2. 2-D clinostat design and growth chamber settings

A total of 12 seedlings were transferred to our modified Bio-World GA-7 “Magenta” growth vessels. We simulated microgravity for the experimental kale by means of the 2-D clinostats set to 10 rpm, which we purchased through COSE (insert citation from paper A). Both the experimental and control plants were grown in a Conviron ATC26 growth chamber for 6 weeks. The temperature of the growth chamber was kept at 22.5°C, with CO_2_ levels reported to be between .06-.08%

### 2.3. Extraction for HPTLC and NMR analysis

Leaf samples were harvested 6 weeks post-planting of seedlings. and subsequently freeze dried for analysis. Samples were ground using IKA sterile grinder at 4k rpm, Samples were weighed in 100mg amounts and were added to 5.00 mL of HPLC grade methanol: water 70:30 v/v. Samples were sonicated for 30 minutes at 30C. Samples were centrifuged and filtered for .45 um filters

### 2.4. NMR experiments

All NMR experiments were performed at 303 K on a Bruker Avance II 400 MHz spectrometer equipped with a triple resonance probe (^1^H, ^13^C, and ^31^P). 1-D data were visualized and processed using the Bruker software Topspin version 4.0.7. All experiments were run using 128 scans, a sweep width of 20ppm and a carrier position set to 6.175ppm and the TD was set to 65K. Samples were run in biological triplicates. Topspin data processing of NMR results

All experiment files were calibrated using a DSS peak as an internal standard. The peaks were picked using the manual semi-automated peak picking function. All phasing and peak picking was done specific to the aromatic region (6.5-8.5ppm). Pick picking minimum relative intensity was set to 2.5. This was done to ensure there was no peak picking of the peaks which were too close to the baseline of the spectrum. This allowed us to confidently analyze exclusively the prominent peaks. While the peak picking rendered near-perfect chemical shift matches between the biological triplicates, there were some minor deviations in the intensities of the peaks; this is to be anticipated as they are biological replicates. The deviation in the peak intensities can be noted in the standard deviation within figure 3.1.

**Figure 3.1.**
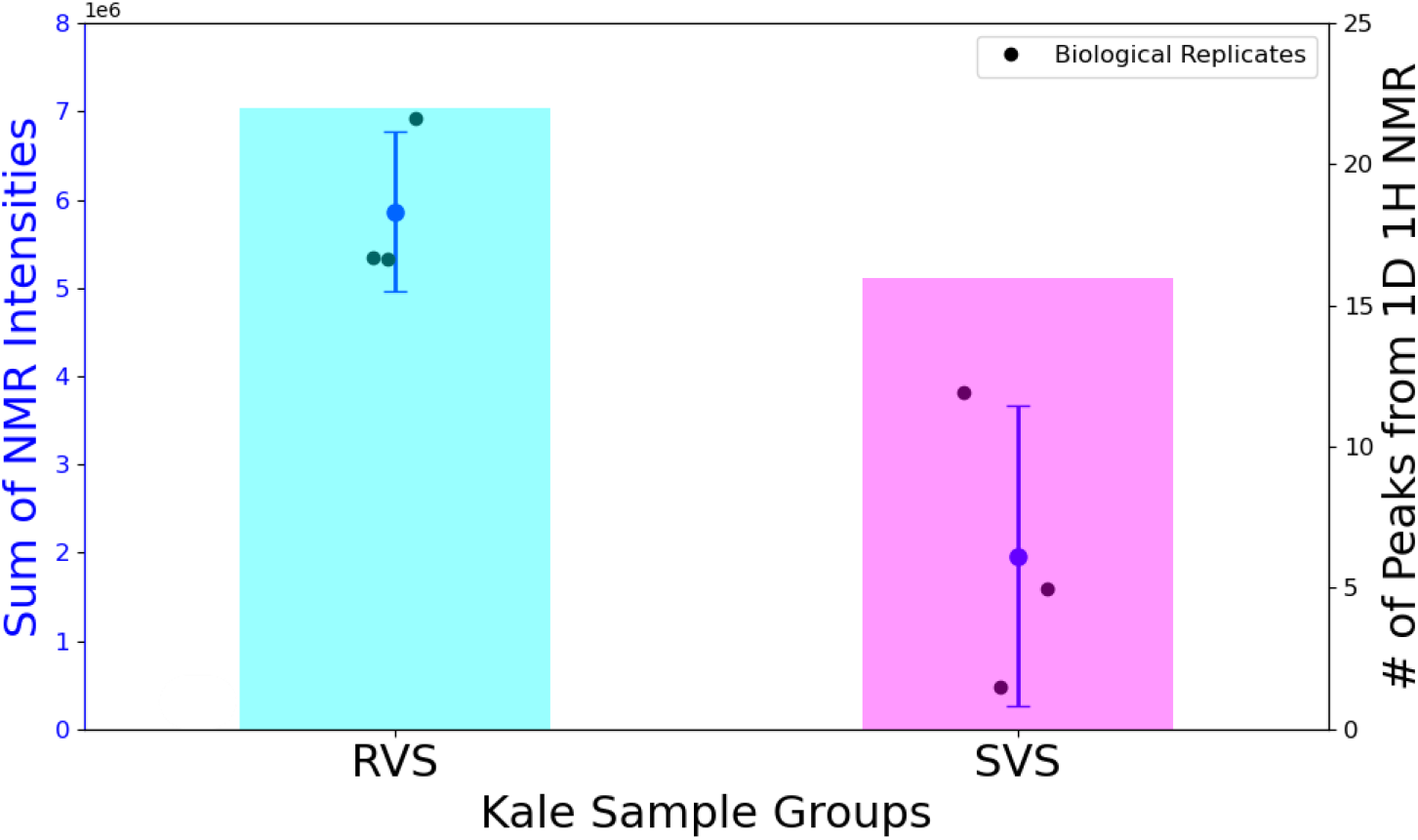
Bar charts comparing the differences in peaks present within the region of 6.5-8.5 ppm of the 1D proton NMR spectra from kale samples grown in different growth conditions: (Cyan) RH Rotating Horizontal: simulated microgravity and (Magenta) SV Stationary Vertical: gravity.

### 2.6. HPTLC experiment

The experimental and control kale samples extracted were concentrated and were resuspended in pure methanol. Our analysis was performed on silica gel 60 F254 glass HPTLC plates 20×10cm. We used several HPTLC instrumented which were manufactured by CAMAG (Muttenz, Switzerland). Experiments were accomplished with the following instruments: Automatic TLC Sampler (ATS 4), Automatic Developing Chamber (ADC 2), Derivatizer, Visualizer 2, TLC Scanner 4 and the visionCATS software version 3.1. Our kale extracts were spotted on the plates at a rate of 150 nL/s as 8 mm bands, 13 mm apart and 8 mm from the lower edge of the plate, with the first track 24 mm from the edge of the plate. We adjusted the plate activity by maintaining a condition of 33% relative humidity (r.H.) with a saturated aqueous solution of magnesium chloride (MgCl2) for 10 minutes. After adjusting the plates, the developing occurred with chamber saturation (20 min, with saturation pad), or without saturation, to a distance of 70mm from the lower edge of the plate, and then dried for 5 min. We derivatized our plates with Natural Product (NP) reagent. However, before derivatization we heated the plates at 100 °C for 3 min. Plates were afterwards dipped in 170.00 mL of chilled methanol with 20.0 mL acetic acid and 10.0 mL of sulfuric acid and 1.0 mL of anisaldehyde (p-methoxy benzaldehyde). Subsequently, the plate was heated for 3 minutes at 100°C. For all analyses, images of plates were captured in UV wavelengths of 254nm, 366nm broadband) and white light in reflection/transmission (RT).

## 3. Results and Discussion

### 3.1. NMR experiment shows flavonoid content differences

Our NMR data (Figures 3.1 and 3.2) readily signals the potential impact on flavonoid content due to the stressors of simulated microgravity of kale. This research used a 1-D proton probing NMR experiment to gauge the effects of the different conditions of the kale vegetables. This very approach has been implemented in recent publications ^37-38^ for untargeted and targeted metabolomic profiling of various samples. The approach is rooted in the understanding that flavonoids are known to appear in the 6-8ppm range of a 1D proton spectrum given their aromatic properties ^39^. The presence of peaks indicates the potential for those peaks to pertain to flavonoids or similar metabolite structures. Therefore, it was reasonable to postulate that spectra which comparatively contained quantities of peaks in their 6-8ppm ranges, inherently possessed more varieties of flavonoids or metabolites similar in structure. We note that the kale grown in simulated microgravity conditions possessed more peaks, and elevated peak intensities than gravity grown kale (approximately 30% more). This hints that the simulated microgravity conditions generated a greater variety of flavonoids or flavonoid-like structures than gravity. This aligns well with our hypothesis that stressful growth conditions may have an impact on the metabolomic content of the kale.

### 3.3. HPTLC plates show differences in flavonoid profiles

Our HPTLC experiment showed metabolites, in the form of bands, which pertained to different polarities that were extracted in methanol. We applied these extracts to plates, in triplicates, that were ran in three different polarity mobile phases: medium polar, high polar and low polar (Table 3.1).

**Table 3.1.**
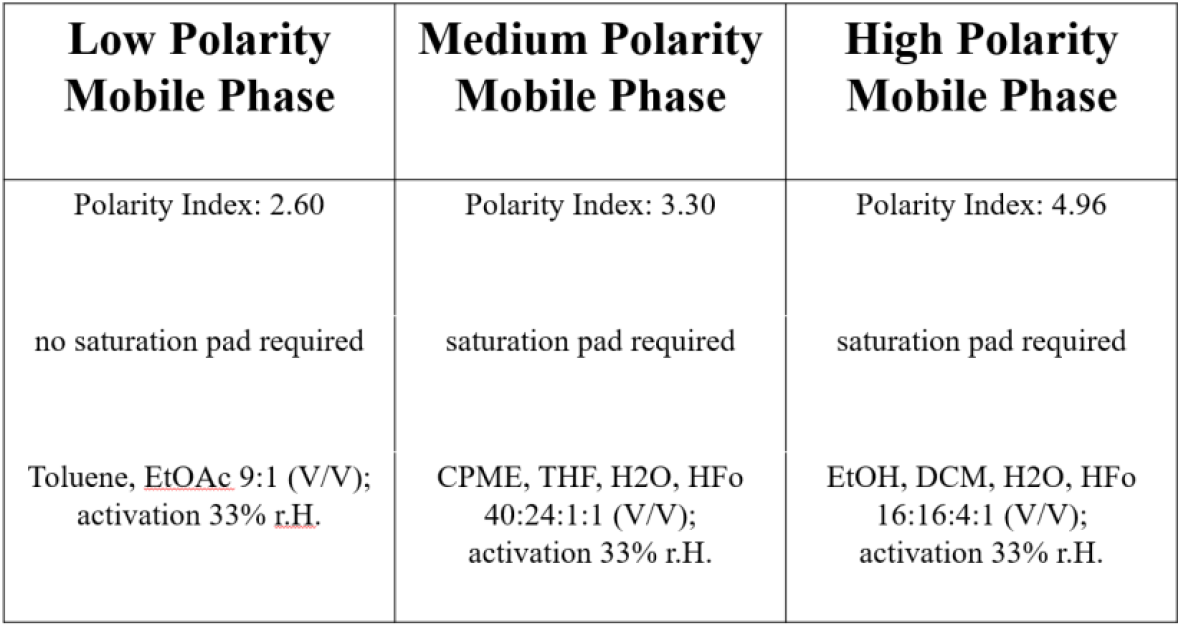
The mobile phases constituting the three polarities used to study kale extracts, and their use of saturation pad parameters.

| **Low Polarity**<br>**Mobile Phase** | **Medium Polarity**<br>**Mobile Phase** | **High Polarity**<br>**Mobile Phase** |
| --- | --- | --- |
| Polarity Index: 2.60 | Polarity Index: 3.30 | Polarity Index: 4.96 |
| no saturation pad required | saturation pad required | saturation pad required |
| Toluene, EtOAc 9:1 (V/V);<br>activation 33% r.H. | CPME, THF, H<sub>2</sub>O, HFo<br>40:24:1:1 (V/V);<br>activation 33% r.H. | EtOH, DCM, H<sub>2</sub>O, HFo<br>16:16:4:1 (V/V);<br>activation 33% r.H. |

These solvent systems were selected based on prior literature ^36^. When contrasting the plates rendered from the three polarities of mobile phases, we note that medium polar possessed 3 (control) and 8 (experimental) number of bands, high polar had 4 (control) and 6 (experimental) number of bands and low polar contained 3 (control) and 3 (experimental) number of bands. These results suggest that a greater number of metabolites extracted in this solvent system are of non-polar polarity. With standards of Catechin, Rutin, Epicatechin, Quercetin, Isoquercetin and Hyperoside in hand (Figure 3.3), we were also capable of confirming their lack of presence in our kale extracts. Simultaneously, our results indicate that in the cases of medium (Figure 3.4) and high polar (Figure 3.6) mobile phases the number of clearly visible bands is greater in the experimental kale samples. These plates have been labeled with annotations of RHSV or SHSV; such annotations are indicative of simulated microgravity or gravity conditions, respectively.

**Figure 3.2.**
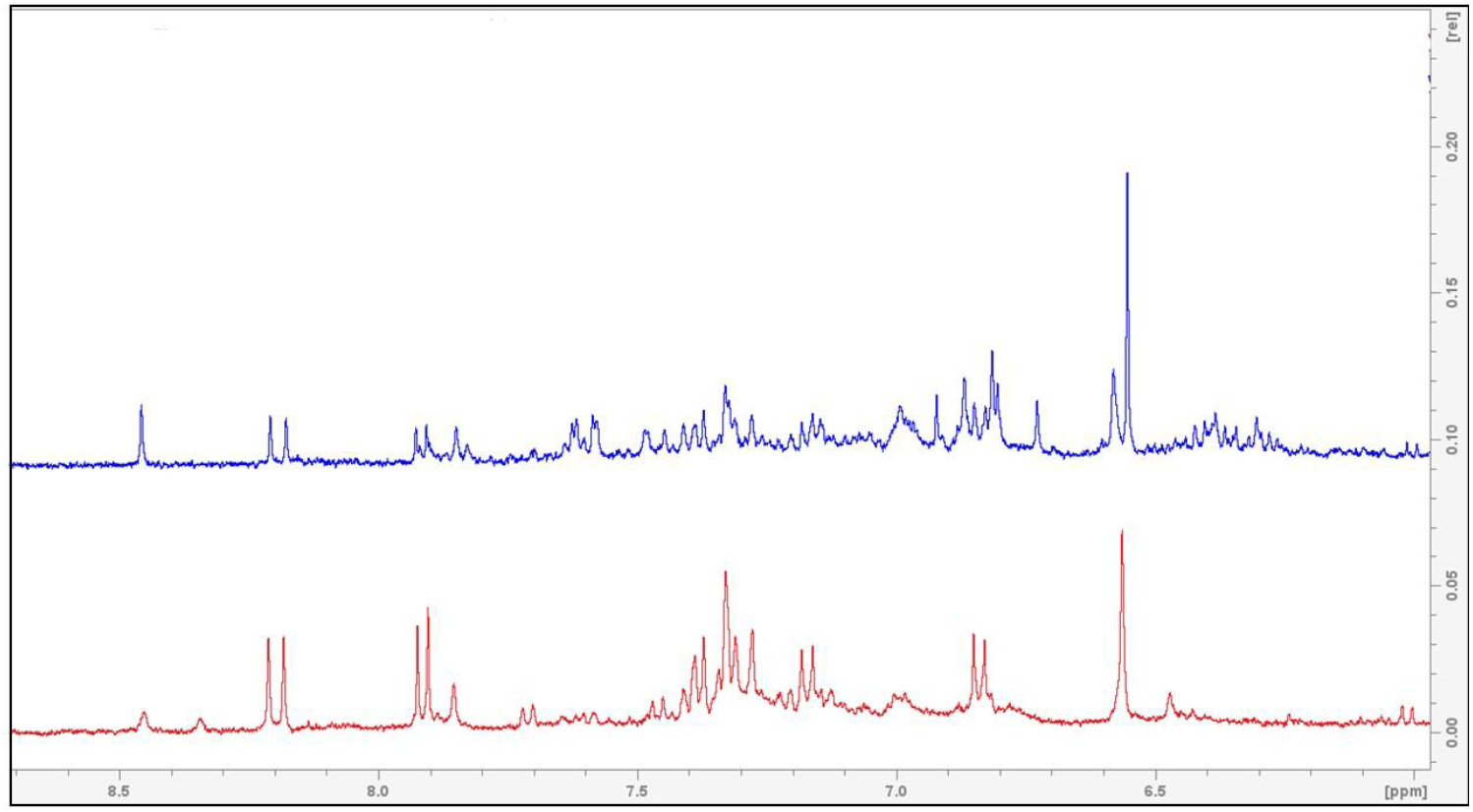
Stacked 1-D ^1^H NMR spectra taken from Topspin version 4.0.7 shows the comparison of the NMR profiles in the aromatic region of 6.5-8.5 ppm from kale samples grown in different growth conditions: (Blue) RH Rotating Horizontal: simulated microgravity and (Red) SV Stationary Vertical: gravity.

**Figure 3.2.**
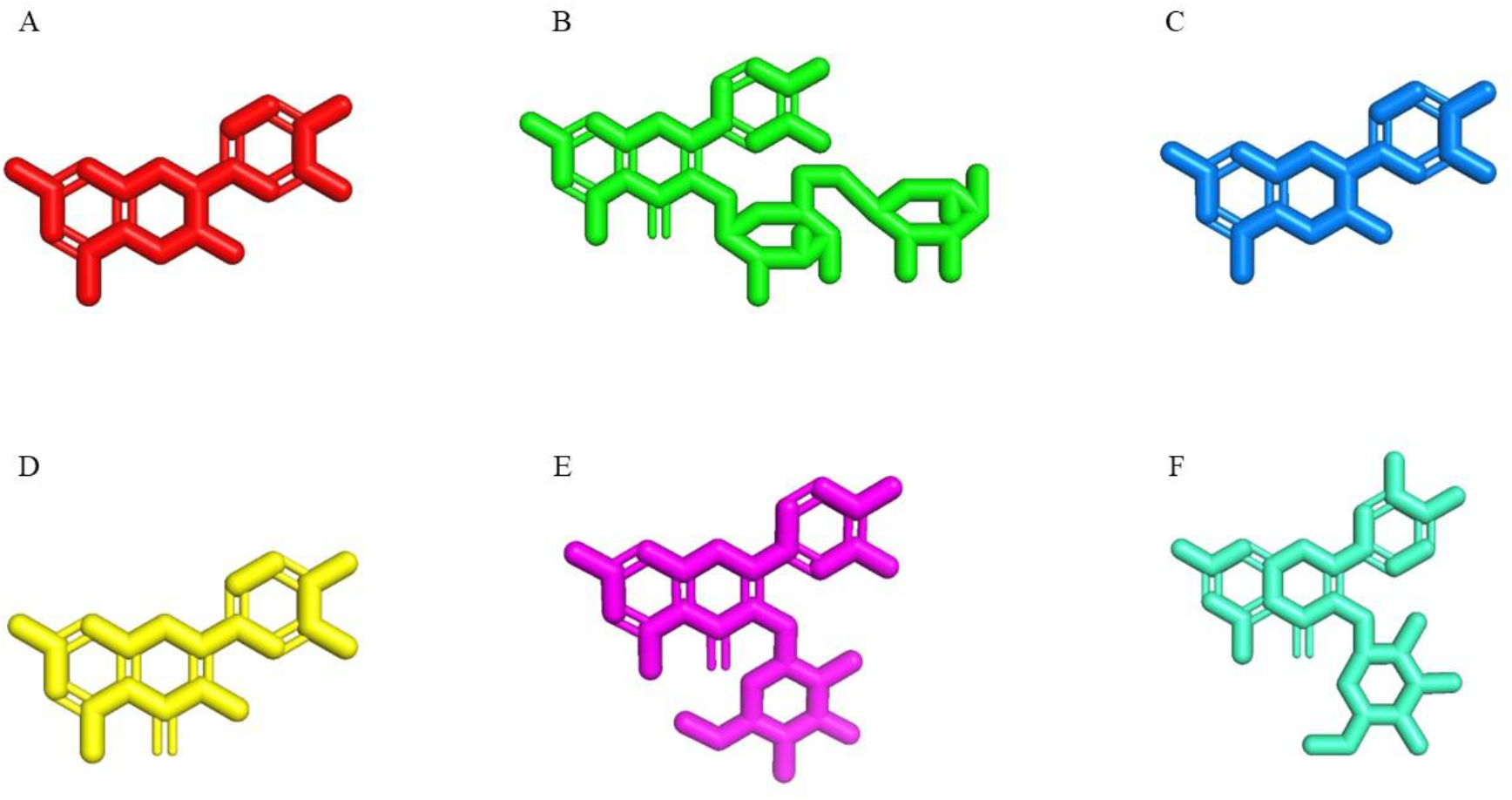
The 3-D structures (PyMOL generated from Chemdraw exports) and compound names of standards used in HPTLC analysis. (A) Catechin, (B) Rutin, (C) Epicatechin, (D) Quercetin, (E) Isoquercetin and (F) Hyperoside.

**Figure 3.4.**
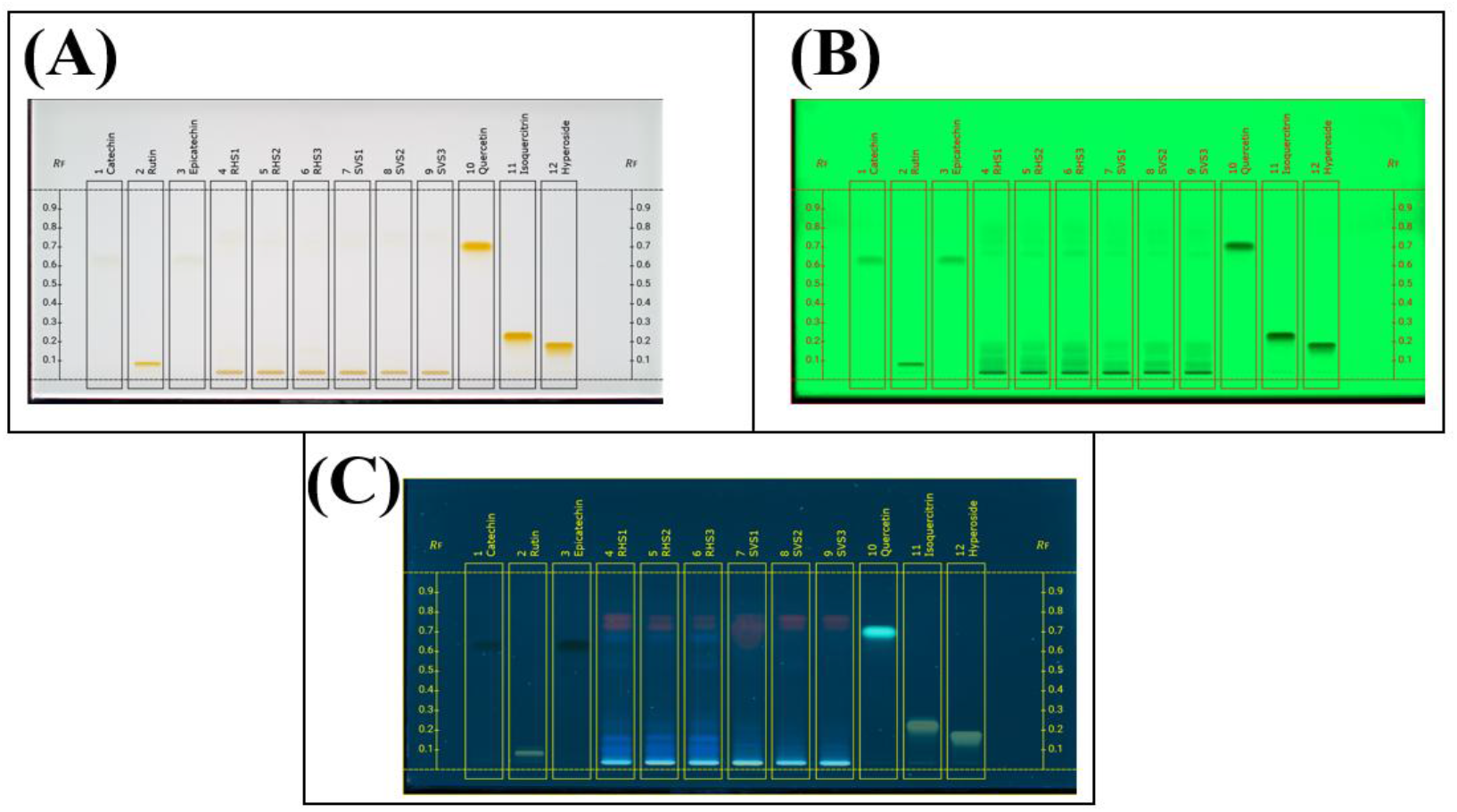
Medium polar plate, taken under **(a)** white light, **(b)** 254nm and **(c)** 366nm which shows the bands extracted in methanol that are of a more semi-polar nature. Plate was run using CPME, THF, H2O, HFo 40:24:1:1 (V/V); activation 33% r.H. (10 min); saturation (20 min, saturation pad).

We report that the low polarity (Figure 3.5) mobile phase renders equal amounts of clearly visible bands in the experimental and control kale (Table 3.3).

**Table 3_3.** The (approx.) number of bands present per mobile phase for control and experimental results HPTLC results.

| **Experimental or control kale** | **Low Polarity Mobile Phase (approx.) number of bands** | **Medium Polarity Mobile Phase (approx.) number of bands** | **High Polarity Mobile Phase (approx.) number of bands** |
| --- | --- | --- | --- |
| Control kale | 3 | 3 | 4 |
| Experimental kale | 3 | 8 | 6 |

**Figure 3.5.**
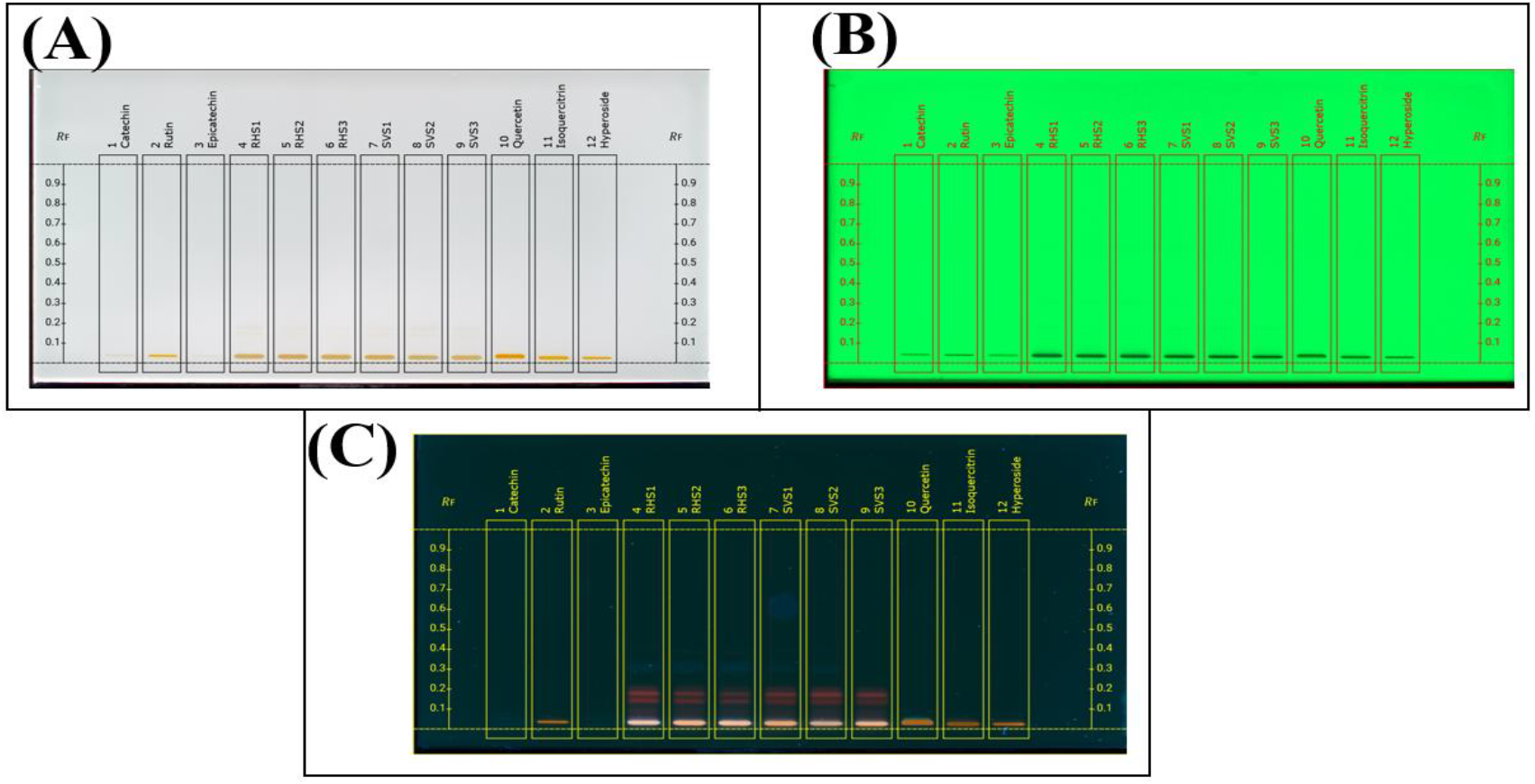
Low polar plate taken under**(a)** white light, **(b)** 254nm and **(c)** 366nm which shows the bands extracted in methanol that are of a more polar nature. Plate was run using Toluene, EtOAc 9:1 (V/V); activation 33% r.H. (10 min); no saturation pad.

**Figure 3.6.**
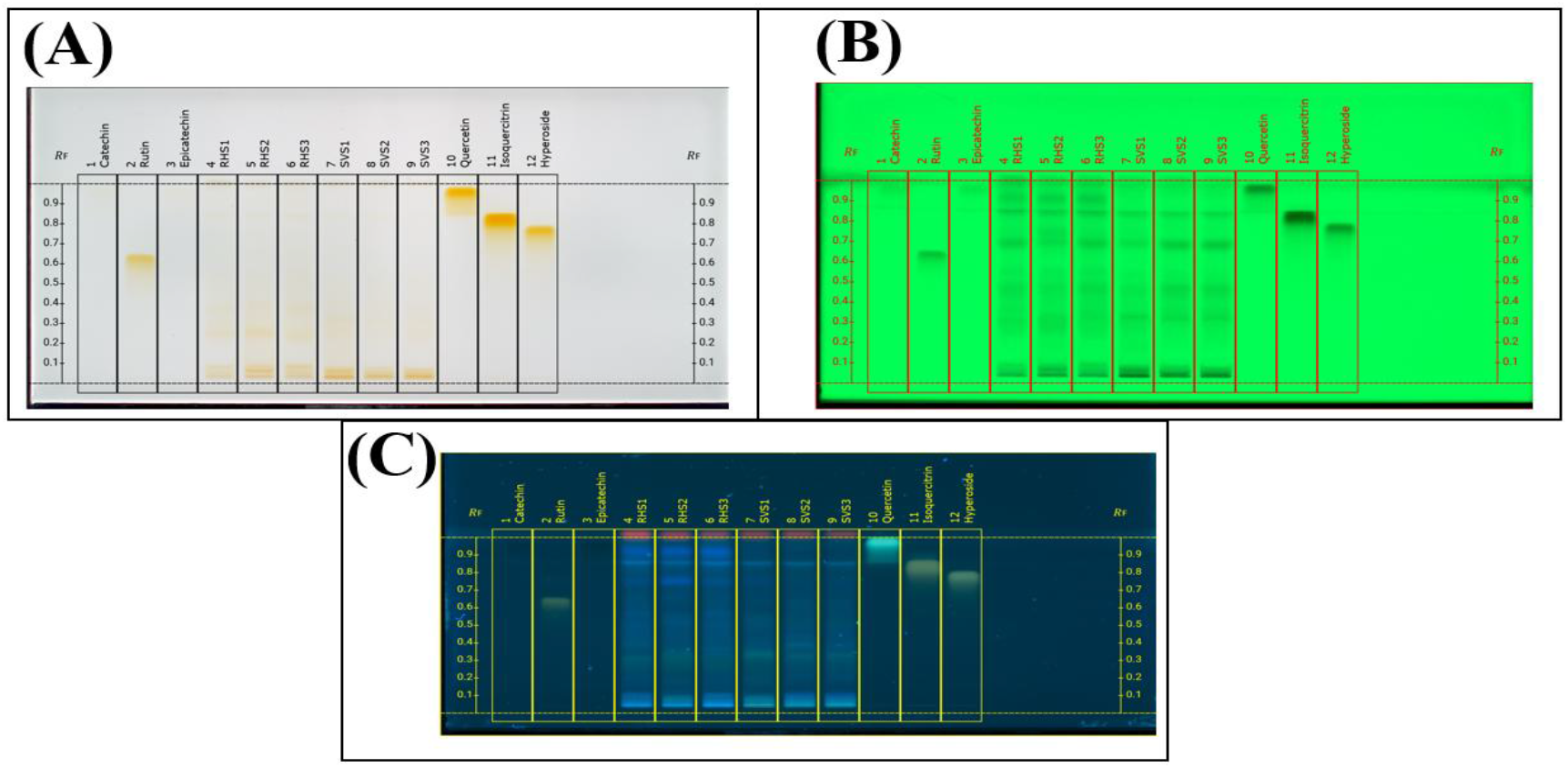
High polar plate taken under **(a)** white light, **(b)** 254nm and **(c)** 366nm which shows the bands extracted in methanol that are of a more non-polar nature. Plate was run using EtOH, DCM, H2O, HFo 16:16:4:1 (V/V); activation 33% r.H. (10 min); saturation (20 min, saturation pad).

In conclusion, this research demonstrates that our kale which was grown in simulated microgravity conditions (experimental) exhibits differences in flavonoids and similar metabolite structure profiles compared to our kale grown in gravity conditions (control). The NMR analysis highlighted a greater amount of peaks in the aromatic region (approximately 30% more), and this signals differences in flavonoid content. Our HPTLC plates co-confirmed the conclusion that the experimental kale produced a higher number of bands in the medium and high polarity mobile phases when compared with the control kale. These results substantiate our hypothesis: the effects of simulated microgravity stress may stimulate the kale’s production of flavonoids and similar metabolite structures. While our work is preliminary, it serves as a meaningful stepping for future studies. Such studies would incorporate longer growth periods of the kale, and investigations into the effects of elevating CO_2_ levels. Yet, our work at hand underscores the potential outcomes of growing flavonoid-rich kale in space to support the health of those in space flight while on spacecrafts.

## Acknowledgements

The authors acknowledge support from NASA (Grant # 80NSSC22K0869).

## Author Contributions

**Rita Dill**: data curation (lead); formal analysis (equal); investigation (equal); methodology (lead); software (equal); validation (equal); visualization (equal); writing—original draft (lead); writing—review and editing (equal).

**Yanru Li**: data curation (lead); formal analysis (equal); investigation (equal); methodology (lead); software (equal); validation (equal); visualization (equal); writing—original draft (lead); writing—review and editing (equal).

**Abdullahi Iro**: data curation (equal); formal analysis (supporting); investigation (supporting); methodology (supporting); software (supporting); validation (supporting); visualization (supporting); writing—original draft (supporting); writing—review and editing (supporting).

**Anne Osano**: conceptualization (lead); data curation (equal); formal analysis (equal); funding acquisition (lead); investigation (lead); methodology (lead); project administration (lead); resources (lead); software (equal); supervision (lead); validation (lead); visualization (equal); writing—original draft (equal); writing—review and editing (equal).

## Data Availability Statement

The data that support the findings of this study are available through contacting Anne Osano

